# Measurement of solubility product in a model condensate reveals the interplay of small oligomerization and self-association

**DOI:** 10.1101/2024.01.23.576869

**Authors:** Aniruddha Chattaraj, Zeynep Baltaci, Bruce J. Mayer, Leslie M. Loew, Jonathon A. Ditlev

## Abstract

Cellular condensates often consist of 10s to 100s of distinct interacting molecular species. Because of the complexity of these interactions, predicting the point at which they will undergo phase separation into discrete compartments is daunting. Using experiments and computation, we therefore studied a simple model system consisting of 2 proteins, polySH3 and polyPRM, designed for pentavalent heterotypic binding. We tested whether the peak solubility product, the product of dilute phase monomer concentrations, is a predictive parameter for the onset of phase separation. Titrating up equal total concentrations of each component showed that the maximum solubility product does approximately coincide with the threshold for phase separation in both the experiments and models. However, we found that measurements of dilute phase concentration include contributions from small oligomers, not just monomers; therefore, a quantitative comparison of the experiments and models required inclusion of small oligomers in the model analysis. We also examined full phase diagrams where the model results were almost symmetric along the diagonal, but the experimental results were highly asymmetric. This led us to perform dynamic light scattering experiments, where we discovered a weak homotypic interaction for polyPRM; when this was added to the computational model, it was able to recapitulate the experimentally observed asymmetry. Thus, comparing experiments to simulation reveals that the solubility product can be predictive of phase separation, even if small oligomers and low affinity homotypic interactions preclude experimental measurement of monomer concentration.

## Introduction

Biomolecular condensate formation by the biophysical process of phase separation is an important modulator of membrane, cytosolic, and nuclear organization in cells (1–3). Phase separation of specific sets of biomolecules underlies the formation of membraneless compartments (4) with unique biochemical environments that organize specific cellular functions much like their membrane-bound organelle counterparts (5–8). For instance, T cell signaling clusters composed of LAT and binding partners promotes local actin polymerization (9), specific interactions with dynamic actin networks at the T cell immune synapse (10), and Ras activation (11). A recent study also highlights functional coupling of lipid and protein phase separation in T cell activation (12). Likewise, nephrin clusters control local actin polymerization by modulating the dwell-time of the Arp2/3 complex at the membrane (13). Aberrant phase separation of biomolecules is implicated in disease, including cancer and neurological disorders (14, 15).

Extensive theoretical and experimental work have been performed to understand the biophysical principles that control the phase separation of biopolymers, including multivalent proteins and nucleic acids (16–20). Many of these studies used model proteins, protein fragments, nucleic acids, or other biomolecules in simplified *in vitro* biochemical reconstitution assays to investigate phase separation (17, 21–25). Key insights into the role of phase separation in the formation and function of biomolecular condensates, and how aberrant phase transitions may underlie disease, emerged from these studies.

Initially, the thermodynamic understanding of liquid-liquid phase separation of polymers, originally described by Flory, was applied to all instances of biological phase separation (26). In the single component (homotypic) version of the model, concentration of a polymer increases to a saturation concentration, above which the solution will demix into a dense phase and dilute phase. Additional polymers added to the solution will then partition into the dense phase. In this model, the concentration of both the dilute and dense phase will remain constant; only the volume fraction of the dense phase will increase as polymers are added to the solution. However, this self-interacting homotypic model cannot capture the complexities of multi-component biological condensates. Recently, cellular experiments showed that not all cellular condensates follow strict Flory-Huggins phase separation principles derived from simple homotypic systems (27).

Indeed, computational simulations (28) predict that saturation concentrations for multi-component condensates may not always remain fixed. Especially for heterotypic interaction-driven systems, dilute phase concentrations of constituent molecules do not reach a plateau beyond the phase transition threshold. A carefully conducted cellular experiment investigating the effect of increased nucleolar scaffold nucleophosmin 1 (NPM1) concentration demonstrated that, instead of fixed dilute and dense phase concentrations with increasing volume fraction as more NPM1 was expressed, concentrations of dense and dilute phases and the volume fraction all increased as the concentration of NPM1 increased in the nucleolus (29). These results illustrate the need for continued development and improvement of models that can be used to predict and dissect biological phase separation in complex systems.

We have previously proposed the concept of solubility product, the product of monomer concentrations (SP), in explaining the saturation concentration of multi-component heterotypic systems (30, 31). In these modeling studies, when SP reaches a maximal level, the system tends to form large multi-molecular clusters, commonly known as percolation or gelation. Beyond this concentration, SP in the dilute phase gradually decreases at the expense of growing clusters. In other words, the modeling studies predict that the maximal SP may characterize the phase transition point for a multi-component system.

In the current study, we combine quantitative biochemical assays and Langevin dynamics simulations to see if the SP concept could be helpful in analyzing the condensation of a simple heterotypic system. We used a 5X tandem repeat of the second Src Homology 3 domain of Nck (polySH3 hereafter), and a 5X tandem repeat of the Abl tyrosine kinase proline-rich motif (polyPRM hereafter), which were previously used to investigate biological phase separation (16, 17, 32), to quantitatively assess dilute and dense phase concentrations of each component. We found that, although experimental data were broadly consistent with our previous proposal that the solubility product predicts phase separation, we identified several factors that complicate the analysis of experimental systems. Importantly, deviations from the expected, ideal behavior of polySH3 and polyPRM predicted by our model enabled the discovery of additional weak interactions between polyPRM proteins that contribute to the phase behavior of our system.

## Methods and Materials

### Cloning, expression, purification, and labeling of polySH3 and polyPRM proteins

5X repeat Nck SH3 domain 2 (polySH3) and 5X repeat Abl PRM (polyPRM) codon-optimized genes were purchased from GenScript and cloned into a pMal plasmid to generate fusion proteins with an N-terminal Maltose Binding Protein (MBP) and C-terminal His_6_ used for affinity chromatography. Each gene contains a single cysteine residue for maleimide conjugated dye labeling. PolySH3 and polyPRM were expressed in *E. coli* strain BL21 DE3^T1R^ by induction with 1 mM IPTG. Cells were lysed using an Avestin Emulsiflex C3 Cell Homogenizer and lysates were centrifuged at 14,000 g for 30 minutes at 4 °C to clear the lysate. Lysates were passed over Ni-NTA Agarose resin (QIAGEN) to purify polySH3 and polyPRM from the solution. MBP and His_6_ tags were removed using TEV protease for 16 hours at 4 °C. Following TEV protease treatment, solutions containing polySH3 and cleaved MBP- and His_6_-tags were passed over Amylose resin (New England Biolabs) to remove excess MBP. TEV-cleaved polySH3 and polyPRM were further purified using anion exchange chromatography (for polySH3, Source 15Q resin (Cytiva)) or cation exchange chromatography (for polyPRM, Source 15S resin (Cytiva)), and size exclusion chromatography using Superdex 200 Increase resin (Cytiva). PolySH3 was labeled using Alexa Fluor 488 (AF488) C_5_ Maleimide (Molecular Probes). PolyPRM was labeled using Alexa Fluor 647 (AF647) C_2_ Maleimide (Molecular Probes) using the manufacturer protocol.

### *In vitro* phase separation assays

Fluorescence microscopy and dynamic light scattering (DLS) were used to evaluate phase separation of polySH3 and polyPRM. Briefly, purified unlabelled poly-SH3 and AF488-polySH3 and unlabelled polyPRM and AF647-polyPRM at concentrations and percentages corresponding to those listed in the results section were mixed in buffer containing 150 mM NaCl, 50 mM HEPES pH 7.3, 1 mM TCEP, and 1 mg / ml BSA at room temperature in an Eppendorf tube and incubated for 2 hours to allow for the equilibration of phase separation. Samples were mixed well and transferred to either a 384 well glass bottom microscopy plate (Greiner) that was previously blocked with 10 mg / ml BSA for 30 min and washed with sample buffer described above to remove excess BSA or 384 well polystyrene bottom plate (Corning) for fluorescence microscopy or dynamic light scattering, respectively.

### Microscopy

Images of polySH3 and polyPRM containing samples were captured using a Diskovery Scanhead with selective pinhole sizing mounted on a Leica DMi8 microscope base equipped with a Hamamatsu C9100-13 EM-CCD camera, 488 nm and 637 nm Spectral Borealis lasers, an ASI motorized XY stage, and Leica and ASI Piezo Z focus drives with a 40 X 1.3 NA oil objective. Images were acquired using Volocity software.

In our initial assays, we found that the partition coefficient (fluorescence intensity inside condensates / fluorescence intensity outside condensates) was ∼100, meaning that the intensity inside condensates was 100X higher than outside condensates. These data were consistent with previous reports (32). However, upon quantifying the low-intensity dilute phase concentration, we noted that the measured intensities were outside the linear range of fluorescence just above the lower detection limit of the EMCCD camera used for image acquisition. Because dilute phase concentrations are the key parameter for calculating the SP of polySH3 and polyPRM, we performed additional imaging of both experimental samples and control concentration curves. Images used for measuring condensate intensity were captured using low laser power and a short exposure time; these settings were used to capture images for the high concentration curve (Figure S1A). The same sample was then imaged again using high laser power and a longer exposure time to capture images used for calculating the concentration of protein in the dilute phase; these settings were used for capturing images for the low concentration curve (Figure S1B).

### Generation of fluorescence concentration curves

AF488-polySH3 or AF647-polyPRM were serially diluted from initial concentrations of 106 µM for AF488-polySH3 and 128 µM for AF647-polyPRM. Fluorescence images of known AF488-polySH3 and AF647-polyPRM concentrations were captured using identical settings to those used to capture images of phase separation assays. Fluorescence intensity was measured using FIJI. Fluorescence intensity was correlated with protein concentrations to generate standard concentration-fluorescence curves (Figure S1) that were used to calculate polySH3 and polyPRM in the dense and dilute phases of images of phase separation assays.

### Quantitative analysis of microscopy images

All images were analyzed using FIJI. To obtain concentration of condensates, condensates larger than 10 µm were selected and the fluorescence intensity of the central region was measured to avoid confounding effects of the point-spread-function (17, 33, 34). The intensities of condensates across five images from each of three technical repeats were averaged for each experimental condition and concentration curves were used to calculate the concentration of polySH3 and polyPRM inside the condensates, as we’ve previously done for other quantitative biochemical assays (9, 10). To obtain concentration of the dilute phase, samples containing condensates were centrifuged at 700 g for 7 min at RT to pellet condensed protein. Condensate-free supernatant was removed from the sample, imaged, and fluorescence was measured. Fluorescence-concentration curves were used to calculate concentration of polySH3 or polyPRM in the dilute phase.

### Dynamic light scattering

Samples of polySH3 and polyPRM were analyzed by DLS using a DynaPro Plate Reader (Waters Wyatt Technologies). 0 µM, 20 µM, and 80 µM of either polySH3 or polyPRM were prepared, incubated at room temperature for 2 hours, added to wells in a 384 well plate, centrifuged at 100 g for 1.5 min at RT to remove air bubbles, and analyzed using auto-adjusted laser power.

### Computational simulations

We used the SpringSaLaD simulation platform (35) to carry out Langevin dynamics simulations. We previously employed SpringSaLaD (30, 31) to study molecular clustering in the context of phase transition. This type of coarse-grained simulations are routinely used (19, 36, 37) in studying phase transitions of proteins and nucleic acids. In this framework, a biopolymer is modelled as a collection of spherical sites connected by stiff springs. The sites are divided into two types – “binders” and “linkers”.

We modelled poly-SH3 and poly-PRM with 15 sites (Figure 2B). Five SH3 domains are interspersed with 10 linker sites; linear molecular length = 42 nm. Similarly, poly-PRM contains five PRM sites linked by 10 linker sites; linear molecular length = 28 nm. For all sites, radius = 1 nm, diffusion constant = 2 µm^2^ / s. A dissociation constant of Kd = 350 µM (Kon = 10 µM^-1^.s^-1^, Koff = 3500 s^-1^) was used to describe SH3 to PRM binding. Simulation time constants, dt (step size) = 10^-8^ s dt_spring (spring relaxation constant) = 10^-9^ s.

We start the simulations by randomly placing N number of molecules in the 3D simulation volume. We titrate N to increase the molecular concentrations. Once the system reaches steady state, we compute the cluster size distribution across multiple stochastic trials.

We construct the model using the graphical interface of the SpringSaLaD software and then run multiple simulations in parallel using the High Performance Computing facility at UConn Health (https://health.uconn.edu/high-performance-computing/). A typical 20 ms simulation (time step = 10 ns) of a system consisting of 3000 sites (200 molecules) takes about 4 hours on a Xeon processor. The execution time, of course, is very sensitive to the simulation conditions like number of molecules, structural constraints within molecules etc.

## Results

### Experimental quantification of polySH3 and polyPRM in the dense and dilute phases

We used a 5X repeated SH3 domain (SH3 domain number two of Nck) connected by GGS linkers of equal length, referred to here as polySH3 (molecular weight = 43 kDa), and 5X repeated proline rich motif (Proline rich motif from Abl tyrosine kinase) connected by GGS linkers of equal length, referred to here as polyPRM (molecular weight = 12 kDa), experimental system to test the model predictions when evaluating multi-component condensate formation. We calculated the concentration of Alexa Fluor 488 labeled polySH3 (AF488-polySH3) and Alexa Fluor 647 labeled polyPRM (AF647-polyPRM) in dilute and dense phases by comparing measured fluorescence with a standard concentration curve produced from images of increasing concentrations of dye-conjugated polySH3 or polyPRM (Figure S1) (33, 34). Because we doped labeled protein at 0.25 % to 5 % depending on the experiment, we divided the calculated concentration of AF labeled protein by the percent labeling to calculate the total concentration of protein in the dilute phase. Protein concentrations in solution were verified by measuring the A280 and A488 (AF488-polySH3) or A647 (AF647-polyPRM) absorbance and calculating the concentration using the extinction coefficient of each protein or dye.

To detect phase separation and determine the concentrations in the dilute and dense phases, we performed quantitative fluorescence measurements on microscope images. Images from titrations of equimolar concentrations of AF488-polySH3 and AF647-polyPRM are shown in Figure 1A. Consistent with previously published reports (16, 32), we observed phase separation at ∼40 µM polySH3 and polyPRM (Figure 1A). We observed that concentrations of both proteins in dilute and dense phases changed as their total concentrations were increased (Figure 1B, C). Dilute phase concentrations of polySH3 and polyPRM increased as they approached a peak at 50 µM and then decreased at higher total concentrations. The concentrations of polyPRM and polySH3 diverged from each other at 60 µM, after which the dilute phase concentration of polyPRM was significantly below the concentration of polySH3 (Figure 1B). This indicates that polyPRM preferentially partitions into the dense phase at higher concentrations (Figure 1C).

**Figure 1.**
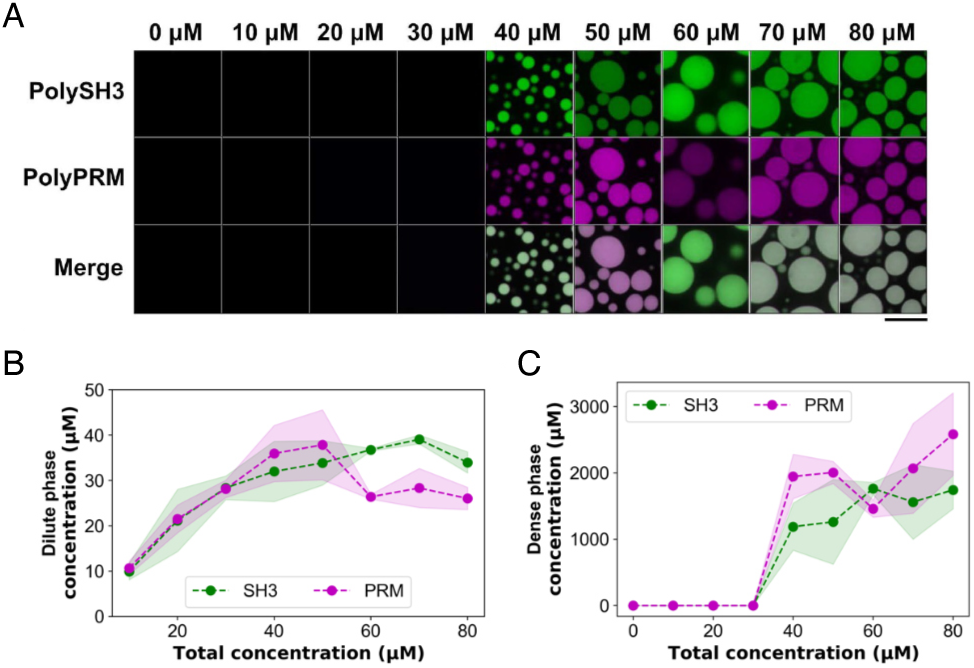
Concentration of polySH3 and polyPRM in dense and dilute phase changes with the increasing total concentration. **A)** Equimolar concentrations of polySH3 (green) and polyPRM (magenta) undergo phase separation above 40 μM each (total concentration of protein = 80 μM). Brightness and contrast for each image were adjusted for visualization and to account for differences in percent labeling. n = 3. Scale bar = 20 μm. **B)** Dilute phase concentrations of polySH3 and polyPRM as a function of total concentration. **C)** Dense phase concentrations of polySH3 and polyPRM as the total concentration increases. In **(B, C)** dilute and dense phase concentrations were calculated from the images obtained for the phase diagram using fluorescence concentration curves (Figure S1A, S1B). The green and magenta lines represent polySH3 and polyPRM, respectively. Shaded regions represent error corresponding to the standard deviation.

### Experimental and computational solubility product trends are qualitatively similar

We used dilute phase concentrations from our experiments where equimolar polySH3 and polyPRM are added in solution to calculate the SP (dilute phase [polySH3] x dilute phase [polyPRM]) and found that the maximum occurred near 40 µM of each component (Figure 2A), coincident with the onset of phase separation (Figure 1A). Beyond the threshold, SP decreases slowly as we increase the total concentrations. (Figure 2A).

To computationally predict the phase behaviour of our model system, we used the SpringSaLaD simulation platform (35) to generate two pentavalent molecules (Figure 2B); one contains five SH3 domains while the other consists of five PRM domains. The lengths of each model molecule approximate the lengths of the actual proteins, with polyPRM being shorter than polySH3. Each SH3 can interact with each PRM with an affinity of 350 µM, derived from the measured binding affinity of the second SH3 domain of Nck and PRM of Abl (16). The experimental solubility product (Figure 2A) is qualitatively captured by our simulations (Figures 2C-E) where the predicted SP, including clusters containing five or fewer molecules, reaches to the maximal level and then decreases (Figure 2C). We chose to include small oligomers ≤ 5 molecules in our solubility product calculations because of previous reports that demonstrate the existence of small oligomers in the dilute phase of a phase separated system (38–40).

To identify phase transitions, we compute the cluster size distribution at each concentration. At a total concentration of 100 μM, molecules were found in small clusters containing less than 15 molecules (Figure 2D, left). However, larger clusters containing more than 20 molecules were observed at a total concentration of 200 μM (Figure 2D, right). The distribution is characterized by a quantity called average cluster occupancy or ACO (Figure S2A, and thoroughly described in (31)). The ACO measures the proportion of molecules residing in large clusters. Since we are titrating up the molecular counts, we normalize ACO by the total molecules (N_total_). When we systematically compute the normalized ACO as a function of total concentration (Figure 2E), it shows a minimum around ∼133 μM. Beyond this point, normalized ACO goes up steadily, indicating that as molecules are being added to the system, they are more and more likely to funnel into large clusters. Thus, the minimum in normalized ACO (ACO / N_total_) serves as the phase transition predictor (31) The histograms produced by the SpringSaLaD simulations above the point of phase separation (Figure 2D) show that a distribution of small oligomers consistently persist in the presence of just 1 or 2 large clusters. These histograms also reveal that below the point of phase separation, the system consists of a distribution of small oligomers, consistent with recent reports (38–40). The oligomer-adjusted SP reaches to a maximal level (Figure 2C) when normalized ACO shows a minimum. Similarly, in the experiment (Figures 1A and 2A) maximal SP coincides with the onset of phase separation. Thus, the notion that maximum SP coincides with the phase transition appears to qualitatively hold for both simulation and experiment.

However, simulated dilute phase concentrations of polySH3 and polyPRM concentrations are similar (Figure S2B), unlike in experiments (Figures 1B, C). Also, we note that a comparison of concentrations at which phase separation occurs are approximately 2-fold higher for simulations than for experiments. This prompted us to further explore both the assumptions underlying our model system to see if we could uncover causes contributing to these quantitative discrepancies.

### SPs of non-equimolar polySH3 and polyPRM concentrations are asymmetrically distributed across the phase diagram

To further explore the utility of the SP parameter, we performed assays from 0 µM to 80 µM at 10µM intervals for the full matrix of combinations of polySH3 or polyPRM. Surprisingly, the experimental phase behavior (Figure 3A) is markedly asymmetric, with a clear bias above the diagonal, indicating that more polyPRM than polySH3 is required for phase separation. When we measured the concentration of polySH3 and polyPRM in the dense and dilute phases, we found that both dilute and dense phase concentrations vary depending on the total concentration of each component used for the assay (Figure S3). These data are consistent with previous observations of measured nucleolar protein concentrations (29). We observe that polyPRM has a greater tendency to partition into condensates; correspondingly, in the 2-phase region, the polyPRM is generally lower in the dilute phase than PolySH3 (Figure S3). This produces an experimental SP pattern that is opposite to the experimental phase diagram (Figure 3B).

**Figure 2.**
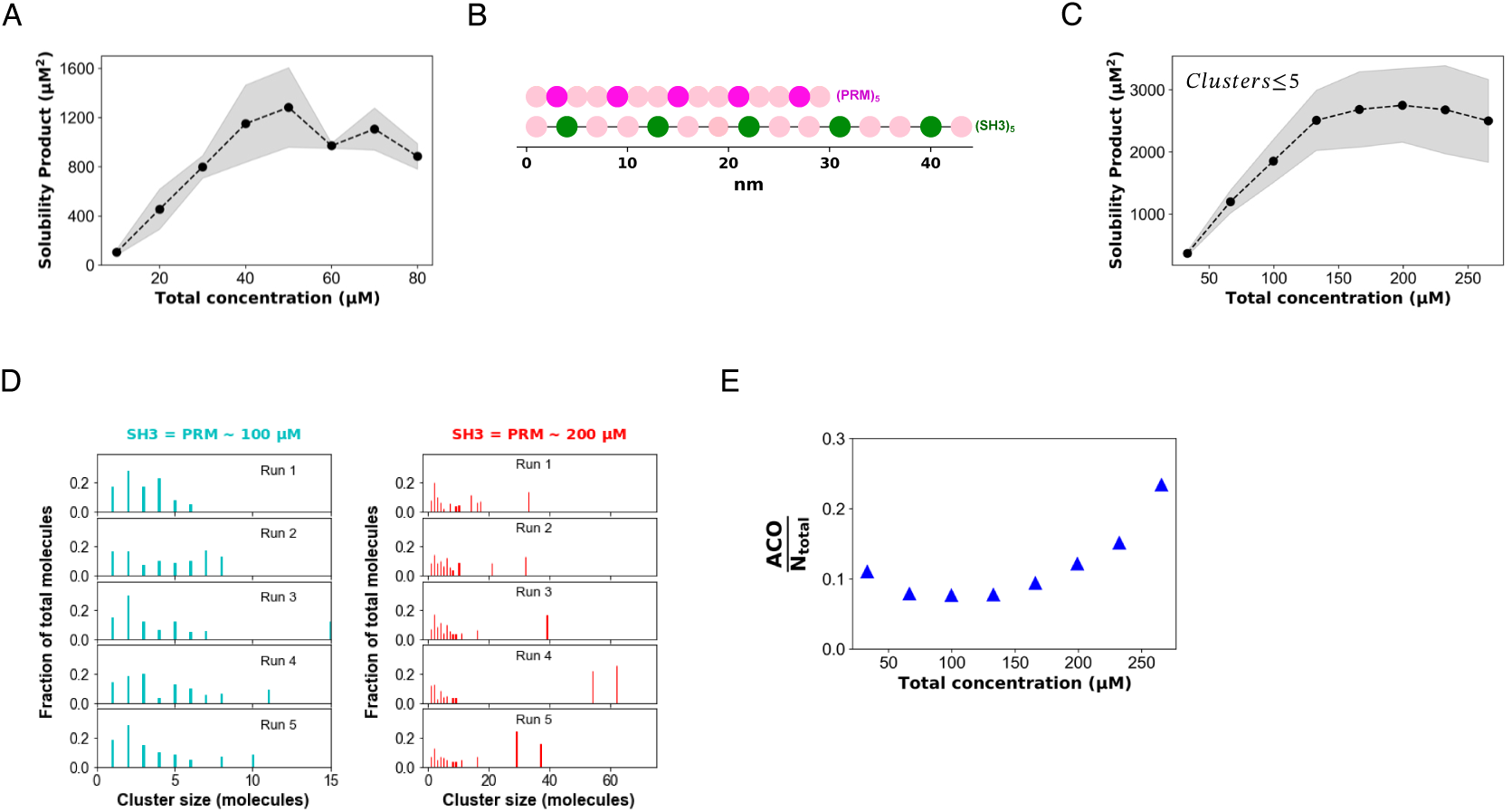
Accounting for small oligomers in our simulated dilute phase results in more accurate quantitative predicted dilute phase concentrations and SP. **A)** Experimentally calculated SP of equimolar polySH3 and polyPRM concentrations. Concentration on X axis is per component, i.e., 20 μM Total Concentration = 20 μM polyPRM - 20 μM polySH3. **B)** SpringSaLaD representation of polySH3 and polyPRM. In polySH3, five (green) SH3 domains are interspersed with 10 linker sites (pink); linear molecular length = 42 nm. Similarly, polyPRM contains five (magenta) binding sites linked by 10 (orange) linker sites; linear molecular length = 28 nm. For all sites, radius = 1 nm, diffusion constant = 2 µm^2^ / s. **C)** Solubility product (SP_dilute_) profile where SP_dilute_ = [Dilute SH3] * [Dilute PRM], derived from simulation results when clusters including up to five molecules are included in dilute phase calculations. **D)** Molecular clusters at 100 µM and 200 µM, respectively. We randomly place N molecules of each type in a 3D reaction volume of 100*100*100 nm^3^. We then titrate up N to increase the molecular concentrations and quantify the cluster size distribution at steady state. For this system, N = [20, 40, 60, 80, 100, 120, 140, 160]. We run 50 stochastic trials for each condition and sample 2 steady state timepoints for each trial. So, each free concentration data point is an average over 100 independent realizations. **E)** Extent of molecular clustering as a function of concentration. Average of the cluster size distribution is referred to as the average cluster occupancy (ACO) which is then divided by the total number of molecules (N_total_) present in the system. In **(A)** and **(C)**, circle represents the mean, and the fluctuation envelope visualizes standard deviations.

**Figure 3.**
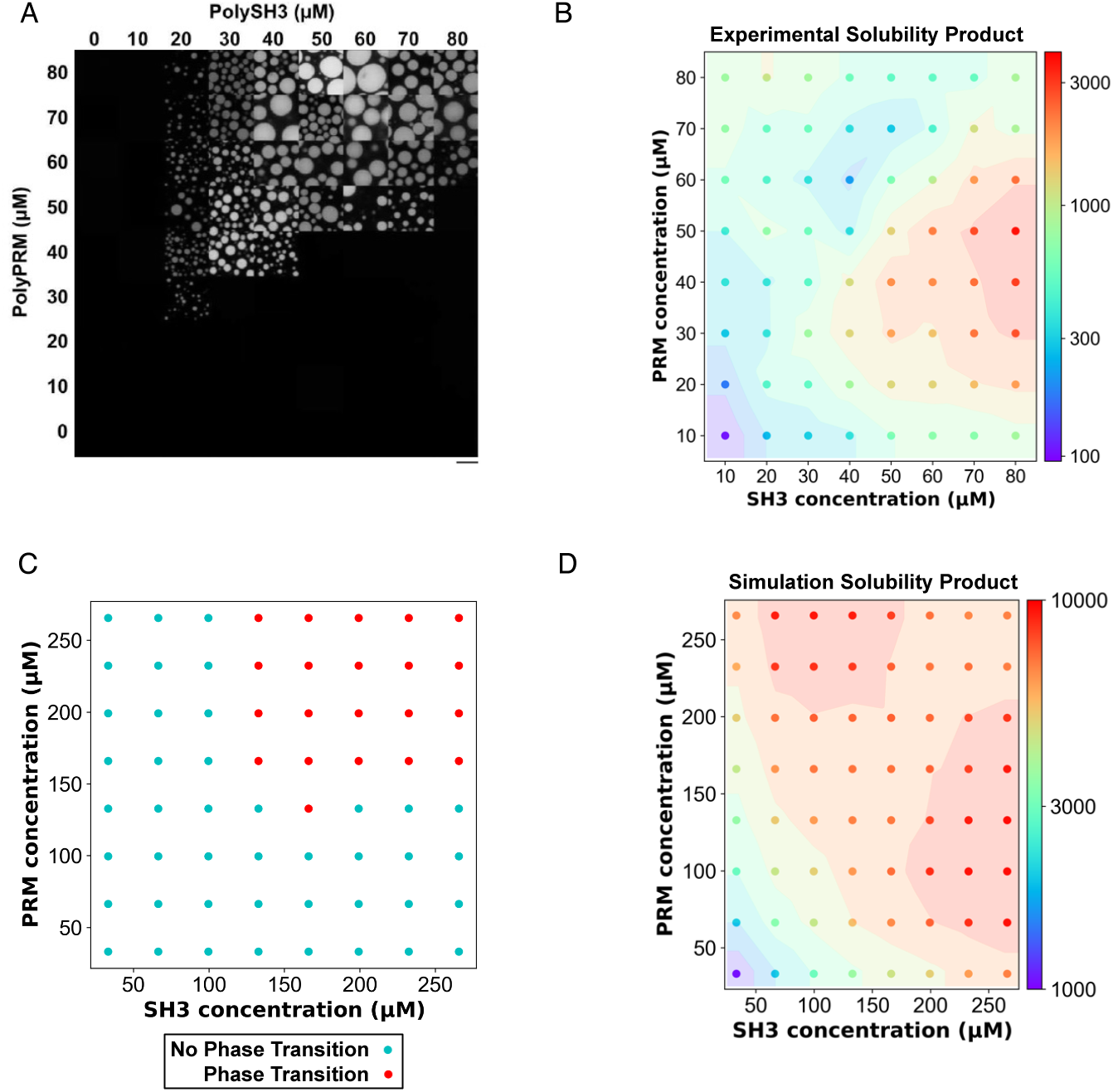
Experimental SPs are asymmetrically distributed across the phase diagram contrary to symmetrically distributed simulated SPs. **A)** Phase diagram obtained using poly SH3 and polyPRM concentrations ranging from 0 μM to 80 μM. n = 3. Scale bar = 20 μm. **B)** Experimental SPs are asymmetrically distributed across concentrations. Color scale indicates the experimentally calculated SP. **C)** Binary classification of phase transition tendency based on the SpringSaLaD simulated cluster size distribution. Red points indicate phase transition while cyan points do not predict phase transition. For the classification, we inspect each steady state timeframe and look for a cluster that contains at least 10% of the total molecules. This thresholding yields a series of “yes” and “no”. If fraction of “yes” > 0.5, then the system has a phase transition tendency. SH3 and PRM counts are 20, 40, 60, 80, 100, 120, 140, 160 and reaction volume is 100*100*100 nm^3^. Number of steady state timeframes = 100. **D)** Solubility product (SP) profile from simulations along the phase diagram where SP = [Dilute polySH3] * [Dilute polyPRM]. As described earlier (Figure 2C), dilute phase includes clusters up to size 5. Number of steady state timeframes = 150.

Concomitantly, we performed simulations across the same range of polySH3 and polyPRM concentrations. In contrast to the experiment (Figure 3B) these resulted in an approximately symmetric pattern of condensate formation on either side of the diagonal (Figure 3C) and a symmetric distribution of SPs calculated using monomer and small oligomer concentrations (Figure 3D). These differences between our simulation and experimental results indicate that the computational model needs to account for additional interactions that might be present in the experimental system. Therefore, we experimentally investigated potential causes of the asymmetry observed in our system.

### PolyPRM self-association contributes to the asymmetry of SPs across polySH3 and polyPRM concentrations

In our simulations, only heterotypic binding of the 2 pentavalent species (SH3 + PRM) is considered. We reasoned that if polySH3 or polyPRM self-associated (homotypic interactions), this could generate situations where molecules are unequally funnelled into condensates resulting in asymmetric concentrations in the dense and dilute phases. Using dynamic light scattering (DLS) we probed the propensity for polyPRM and polySH3 to self-associate. Indeed, polyPRM self-associated in the concentration range used in our experiments; when we performed DLS on polyPRM-containing solutions below (20 μM) or above (80 μM) the threshold for heterotypic (polyPRM + polySH3) phase separation, two peaks were observed (Figure 4A). We did not observe macroscopic phase separation for these concentrations of polyPRM (Figure 3A), consistent with polyPRM forming relatively small oligomers below microscopic resolution. In other words, polyPRM displays a tendency to self-associate (homotypic interactions), which is not strong enough to result in phase separation within the concentration range explored here. In contrast, as we increased the concentration of polySH3 in solution from 20 μM to 80 μM, we did not observe any changes in its dispersity indicating that polySH3 did not form small oligomers in solution (Figure 4B). These data indicate that polyPRM can self-associate in solution while polySH3 remains monodisperse.

**Figure 4.**
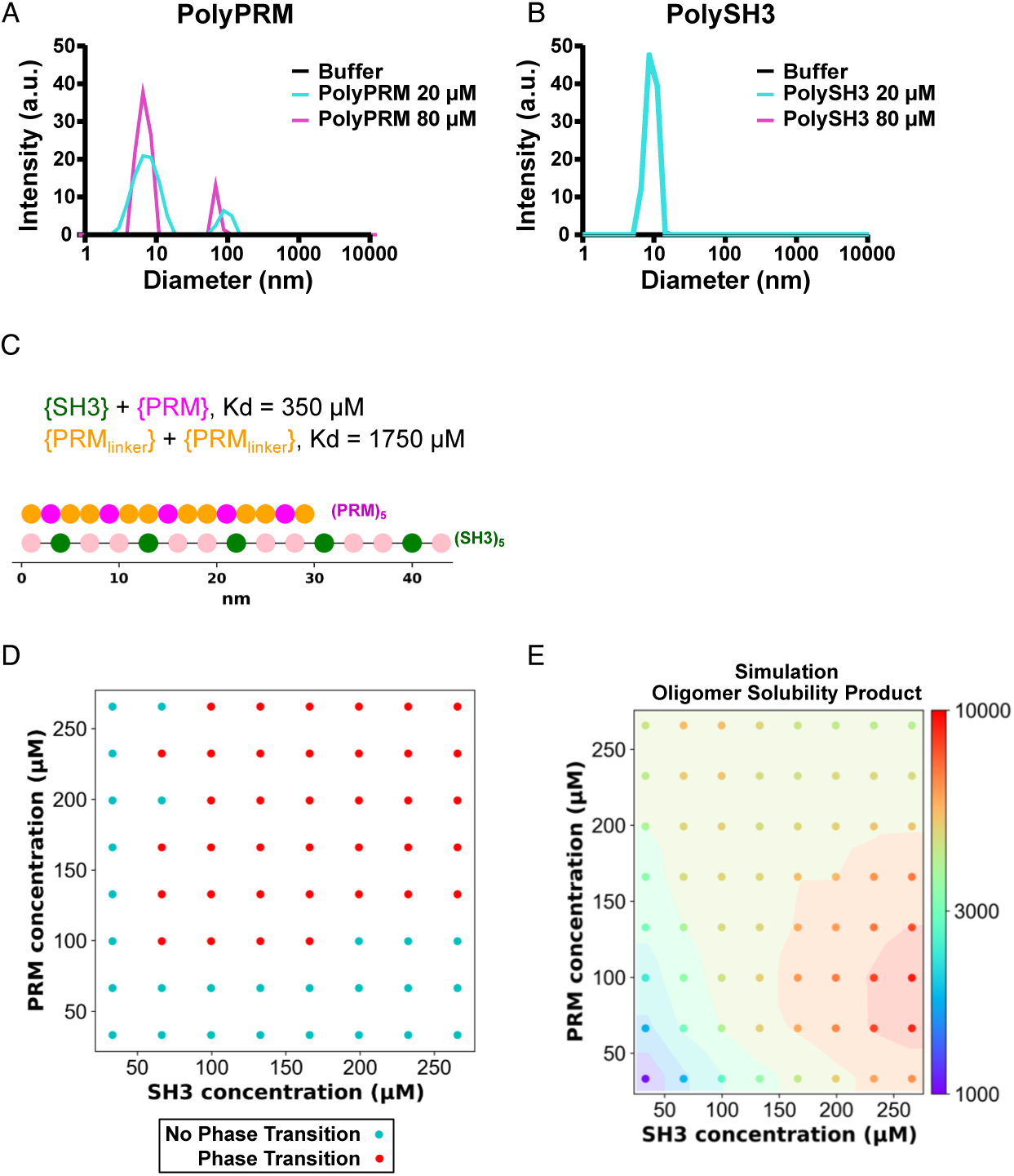
PRM-PRM self-association causes SP to be asymmetrical across the phase diagram. **A)** Dynamic Light Scattering (DLS) analysis of solutions containing polyPRM at concentrations of 0 μM (buffer only, black), 20 μM (cyan), and 80 μM (magenta). **B)** DLS analysis of solutions containing polySH3 at concentrations of 0 μM (buffer only, black), 20 μM (cyan), and 80 μM (magenta). **C)** Illustration of our modified model where PRMs can self-associate. We enabled binding (5x weaker than the canonical SH3-PRM interaction) between linkers of PRM molecules. **D)** Binary classification of phase transition tendency based on the cluster size distribution, as described in Figure 3C. SH3 and PRM counts are 20, 40, 60, 80, 100, 120, 140, 160 and reaction volume is 100*100*100 nm^3^. Number of steady state timeframes = 100. **E)** Simulated SP_oligomer_ (SP that includes small oligomers and homotypic polyPRM interactions) profile across the phase diagram. Number of steady state timeframes = 150.

To model this self-association, we introduced a weak interaction with a binding affinity of 1750 μM (5 times weaker than SH3-PRM binding) between the yellow linker sites in the polyPRM structure (Figure 4C). While these parameters are estimates and not based on experimental data, we felt they would help test the idea that weak self-association of the PolyPRM might allow the model to recapitulate the asymmetric patterns evident in the experiments of Figures 3A and 3B. Indeed, simulations using these parameters showed significantly improved agreement with our experimental data, notably the asymmetry of the phase diagram (Figure 4D vs. Figure 3A). We also observe an asymmetric distribution of SPs across the phase diagram that mirrors our experimentally calculated SPs (Figure 4E vs. Figure 3B). Thus, the inclusion of polyPRM homotypic self-association in simulations recapitulates the asymmetry observed in our experimental data and contributes to more accurate simulations. Also satisfyingly, the simulated phase diagram of Figure 4D is shifted to lower threshold concentrations, compared to Figure 3D, where the additional homotypic interactions had not been considered. Thus, simulations that include the polyPRM homotypic interactions revealed by the DLS measurements are much more quantitatively in line with the experiments for both SP and phase diagrams.

### Extraction of the monomeric SP reveals a symmetric pattern

The correlation between our experimental SP data (Figure 3B) and simulation (Figure 4E) informs us about the validity of our model predictions when we properly account for both small oligomers and the self-association of the polyPRM. We then sought to perform a virtual experiment that examines the behavior of the system based on the monomeric SP. From our simulations, we extract the monomeric concentrations of polySH3 and polyPRM, and compute a monomeric SP (SP_monomer_). Figure 5 reveals that SP_monomer_ is surpisingly symmetric even in the presence of polyPRM self-association. The range at which SP_monomer_ maintains its maximal level decreases in the presence of homotypic interaction, but the pattern remains symmetric around the diagonal line (i.e. the equal concentration line). It is important to note that experimentally resolving monomers using common techniques, including fluorescence microscopy and DLS, is not currently feasible as monomers lie below the resolution limit for these techniques. Although the inherent resolution limit prevents us from experimentally measuring the monomer concentration, our computational model highlights the key interplay of monomers and small oligomers in shaping the composition of the dilute phase.

**Figure 5.**
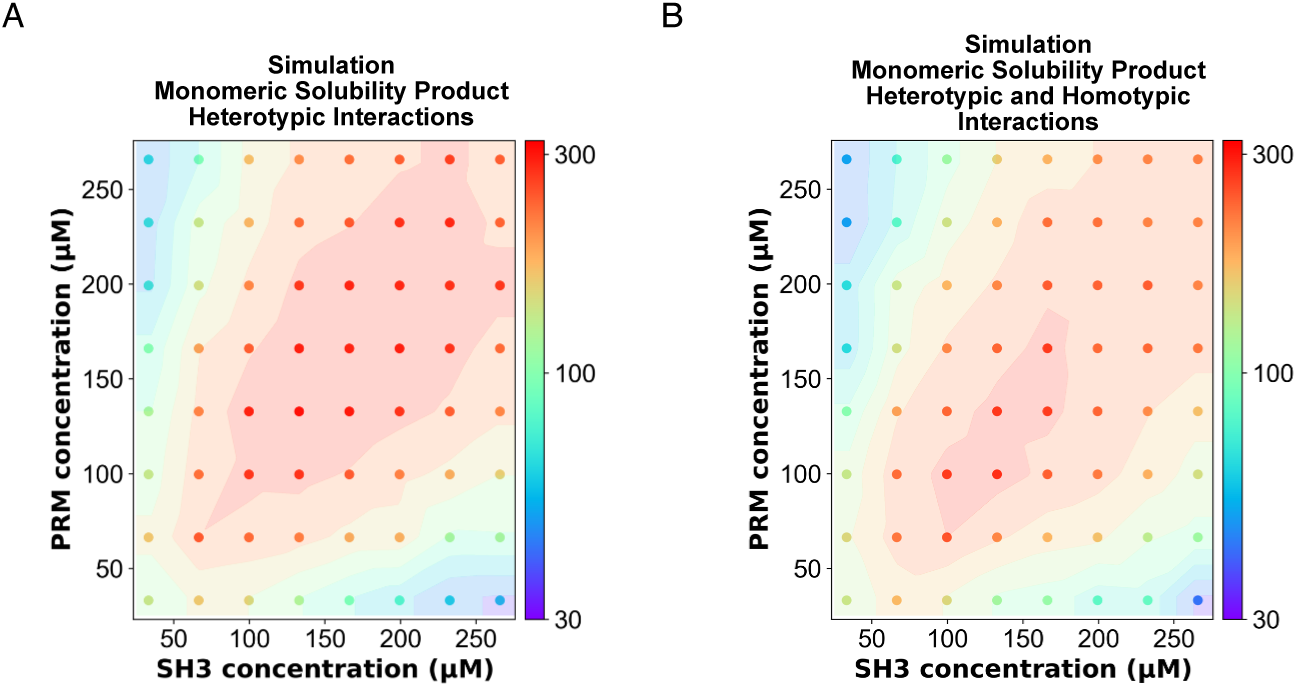
The maximal simulated monomer solubility product occurs at the critical concentrations for phase separation across the phase diagram. **A)** Simulated SP_monomer_ profile across the phase diagram when only heterotypic interactions between polySH3 and polyPRM are modeled. Number of steady state timeframes = 150. **B)** Simulated SP_monomer_ profile across the phase diagram when heterotypic interactions between polySH3 and polyPRM and homotypic interactions between polyPRM and polyPRM are modeled. Number of steady state timeframes = 150.

## Discussion

In homotypic phase separating systems, increasing the concentration of protein or nucleic acid above the critical concentration for phase separation will result in the formation of coexisting dilute and dense phases. Increasing salt, temperature, or pH will alter the critical concentration at which phase separation occurs. In these systems at a given temperature, pH, or salt concentration, each phase will maintain a single concentration as the overall concentration of protein or RNA increases above the critical concentration while the volume fraction of the dense phase will increase. However, biomolecular condensate formation in cells often depends on multivalent binding between multiple heterotypic components. The lack of a threshold saturation concentration for any individual component of a multi-component heterotypic condensate has been problematic for the formulation of a coherent thermodynamic picture of the concentration dependence for phase transition (29). We previously proposed that instead of looking at the individual components, if we consider the stoichiometry-adjusted product of monomeric concentrations (“Solubility product” or SP), this joint quantity converges to a maximal level when the system undergoes a percolation transition (30, 31). In other words, when these multivalent molecules start to form very large clusters, SP of the system saturates, even though monomer concentrations of individual molecular types may continue to increase. Beyond this point, SP gradually goes down as more and more monomers are funnelled into the large clusters. In this current work, we wanted to test our computational predictions using quantitative biochemical assays.

We have previously used spatial Langevin dynamics simulations to show that heterotypic phase separation can be predicted by calculating the SP of heterotypic monomers at equimolar concentrations of model proteins (30, 31). In our initial publication, we found that the SP plateaued at total concentrations that coincided with the formation of mesoscale condensates in our simulations. Using non-spatial simulations, we found that intra-cluster binding, or interactions between molecules that result in the formation of bi-molecular rings is a strong contributor to modeled phase separation (41). More recently, we showed that intra-cluster binding will result in a peak SP at the point where proteins demix into phase separated condensates, followed by a decrease in the SP as the concentration of heterotypic interactors increases (31). In the present study (Figures 1 and 2), experimental results for titrations of equal concentrations of polyPRM and polySH3 are qualitatively similar to simulations that accounted for SH3-PRM interactions and intra-cluster binding. The SP increases with increasing equimolar concentrations and then decreases with increasing polySH3 and polyPRM concentrations above the critical concentration for condensation. Experimentally, the observed decrease in the dilute phase is accompanied by an increase in the dense phase, indicating that more molecules are funneled into the condensates as the total concentration increases.

However, determination of SP across the entire phase diagram revealed a divergence between computational predictions and experiments. We experimentally observed an asymmetrical patterns of SPs and phase separation in our experiments, while our simulations predicted a symmetrical patterns (Figure 3). We hypothesized that this may be caused by self-association interactions between polySH3 or polyPRM. Our DLS analysis of polyPRM in solution experimentally confirmed that polyPRM does self-associate at concentrations at and above 20 μM. Consequently, we included polyPRM self-association in our simulations using a weak binding constant (1750 μM). Accounting for this self-interaction resulted in computational predictions that more accurately predicted experimental results (Figure 4). This is especially relevant because quantitative experiments typically used to study biological phase separation, such as quantitative fluorescence light microscopy, cannot distinguish between monomers and small oligomers that remain in the dilute phase. Therefore, any calculations of dilute phase concentration from fluorescence necessarily include small oligomers as well as monomers. Computational SP that includes small oligomers and homotypic polyPRM interactions (SP_oligomer_) demonstrates asymmetry across the phase diagram, similar to that calculated from experiments (Figures 3B, 4F). Additionally, computationally derived SP_oligomer_ and dilute phase concentrations (Figures 4E, S4A, S4B) showed a similar trend to experimental results (Figures 1B and 2A).

The agreement between experimental and simulated SP_oligomer_ gave us confidence that our computational model accurately described the interactions driving the corresponding experimental system. This allowed us to parse our simulations to obtain the true monomeric SP across the phase diagram. It is through this analysis that the power of our computational model becomes readily apparent and provides mechanistic insight that confirms our previous predictions that the maximal monomeric SP exists at the phase boundary and is therefore predictive of phase separation.

Figure 6 summarizes our working model (hypothesis) unveiled in the current study. For a system undergoing phase separation, there exists a three-state equilibrium. From a macroscopic point of view, there are two phases – dilute and dense that can be observed and measured via experimental techniques. However, within the dilute phase, there is another equilibrium between monomers and small oligomers, which is difficult to detect experimentally. Based on our simulations, we observe monomeric SP achieves a maximum along the diagonal, even if the weak homotypic interactions of the polyPRM are accounted for (Figure 5B). This principle, manifested in the form of a symmetric phase diagram, is obscured in the presence of small oligomers. Not surprisingly, small oligomers affect the off-diagonal elements (where stoichiometric ratio does not equal 1) more than the on-diagonal elements. The composition of small oligomers can be heavily influenced by weak homotypic interactions. These interactions yield a confounding trend in the experimental SP level. Overall, the interplay between quantitative experimentation and computational modeling enables us to recognize a three-component system, monomer-oligomer-condensate, that governs the phase behavior of the model proteins.

**Figure 6.**
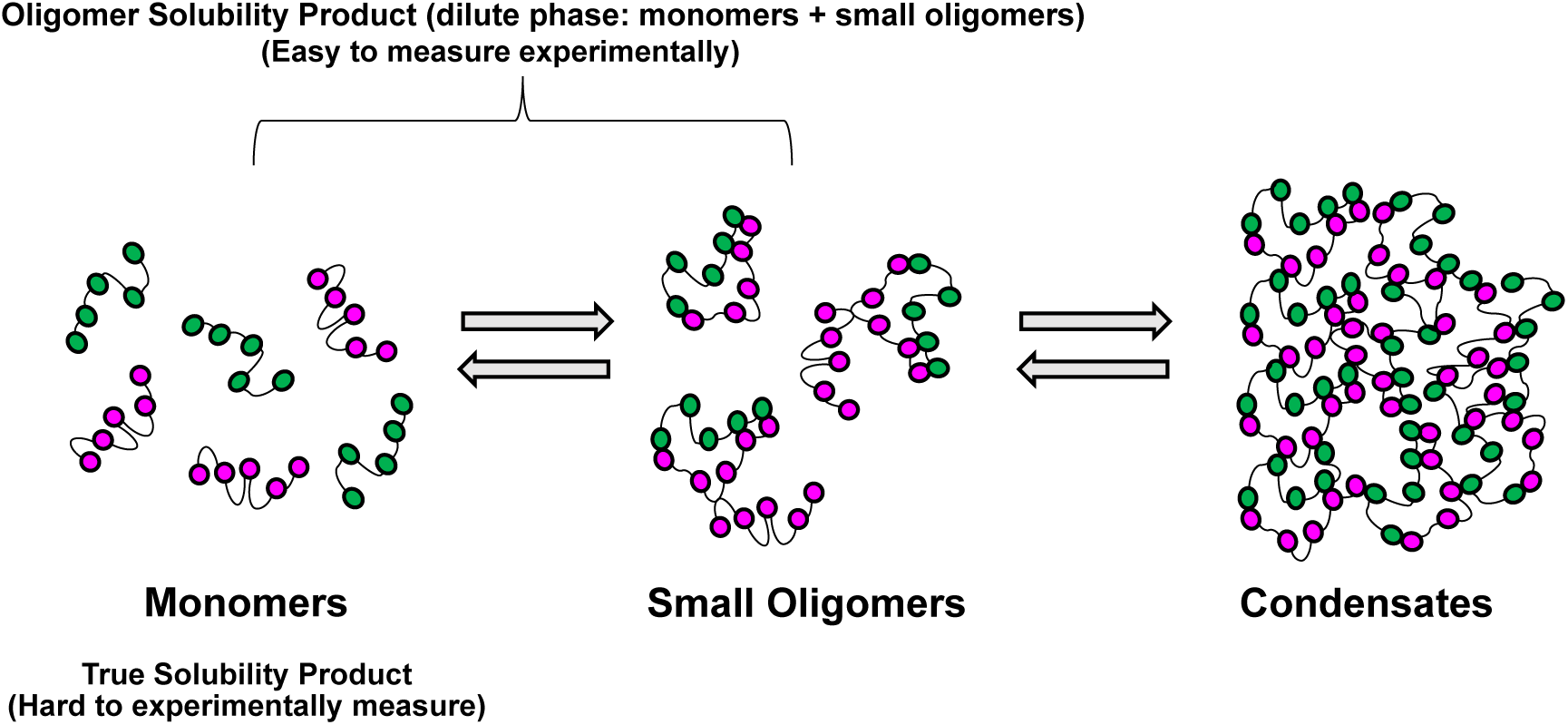
A picture of equilibrium for multicomponent phase separating systems at equilibrium. Experimentally measured dilute phase concentration includes both monomers and small oligomers, allowing us to calculate the SP_oligomer_ across concentrations in the phase diagram. We can also calculate this parameter using computational modeling to determine agreement between experimental and computational systems. When the computational and experimental SP_oligomer_ are in agreement, the model can be parsed for the SP_monomer_. Phase separation occurs at the maximal SP_monomer_.

Despite the presence of homotypic interactions, we used a SP expression that only includes heterotypic interactions. Since the homotypic interactions are not strong enough to initiate a PRM-only phase separation, we feel this approximation is justified. This also allows us to extend the SP idea into more complex multi-component systems. Condensate forming biomolecules can be broadly classified into two categories – “Scaffolds” and “Clients” (4). Scaffolds are required components that drive the phase separation, while clients are auxiliary components that are recruited to the condensates but cannot phase separate on their own. While thinking about the threshold concentration of such complex systems, SP should more strongly depend on the scaffold monomers, and minimally on the client monomers. Overall, mechanistic insights revealed in the current study support the usefulness of the SP idea in explaining the threshold concentration of multi-component biological condensates.

## Acknowledgements

The authors thank Tom Scheuermann, Hyun O. Lee, Joel Watts, and Spencer Freeman for helpful discussions, the SickKids Imaging Facility for technical assistance with microscopy, and the SickKids Structural and Biophysical Core for technical assistance with dynamic light scattering. This work was supported by a Natural Sciences and Engineering Research Council Discovery Grant RGPIN-2022-03274 to J.A.D and NIH grants R24 GM137787 and R01 GM132859 to L.M.L.

## Supplemental Figure Legends

**Figure S1.**
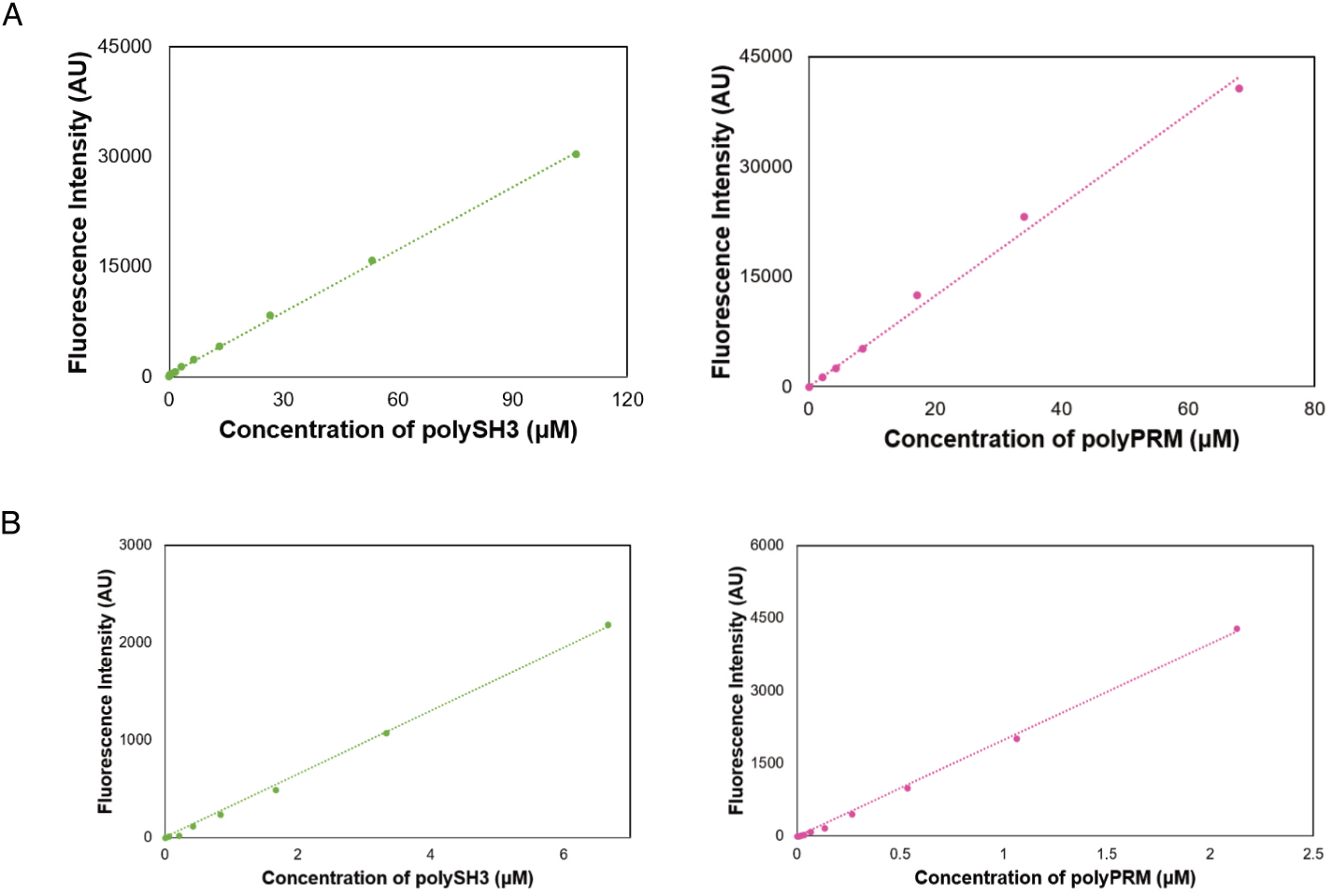
**A)** AF488-polySH3 (left) and AF647-polyPRM (right) dilution curves used for calculating the dense phase concentrations of each protein. **B)** AF488-polySH3 (left) and AF647-polyPRM (right) dilution curves used for calculating the dilute phase concentrations of each protein.

**Figure S2.**
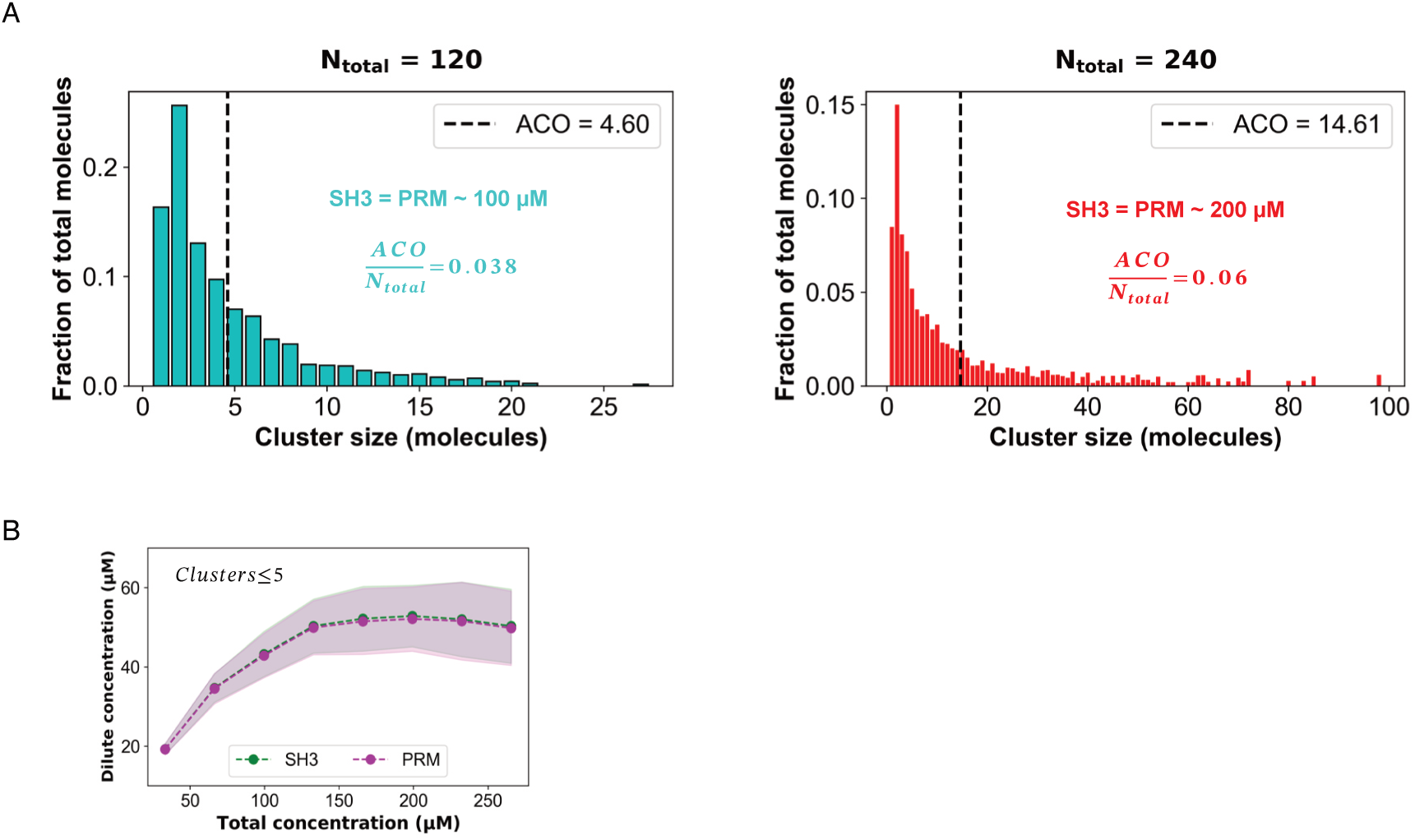
**A)** Simulated fractions of molecules at each size cluster when the concentrations of polySH3 and polyPRM are 120 μM (left) or 240 μM (right). Data used for calculating the ACO in Figure 2E. **B)** Simulated concentrations of polySH3 (green) and polyPRM (magenta) along the concentration diagonal when clusters containing five or fewer molecules are quantified. Simulations only are performed considering heterotypic polySH3 - polyPRM interactions.

**Figure S3.**
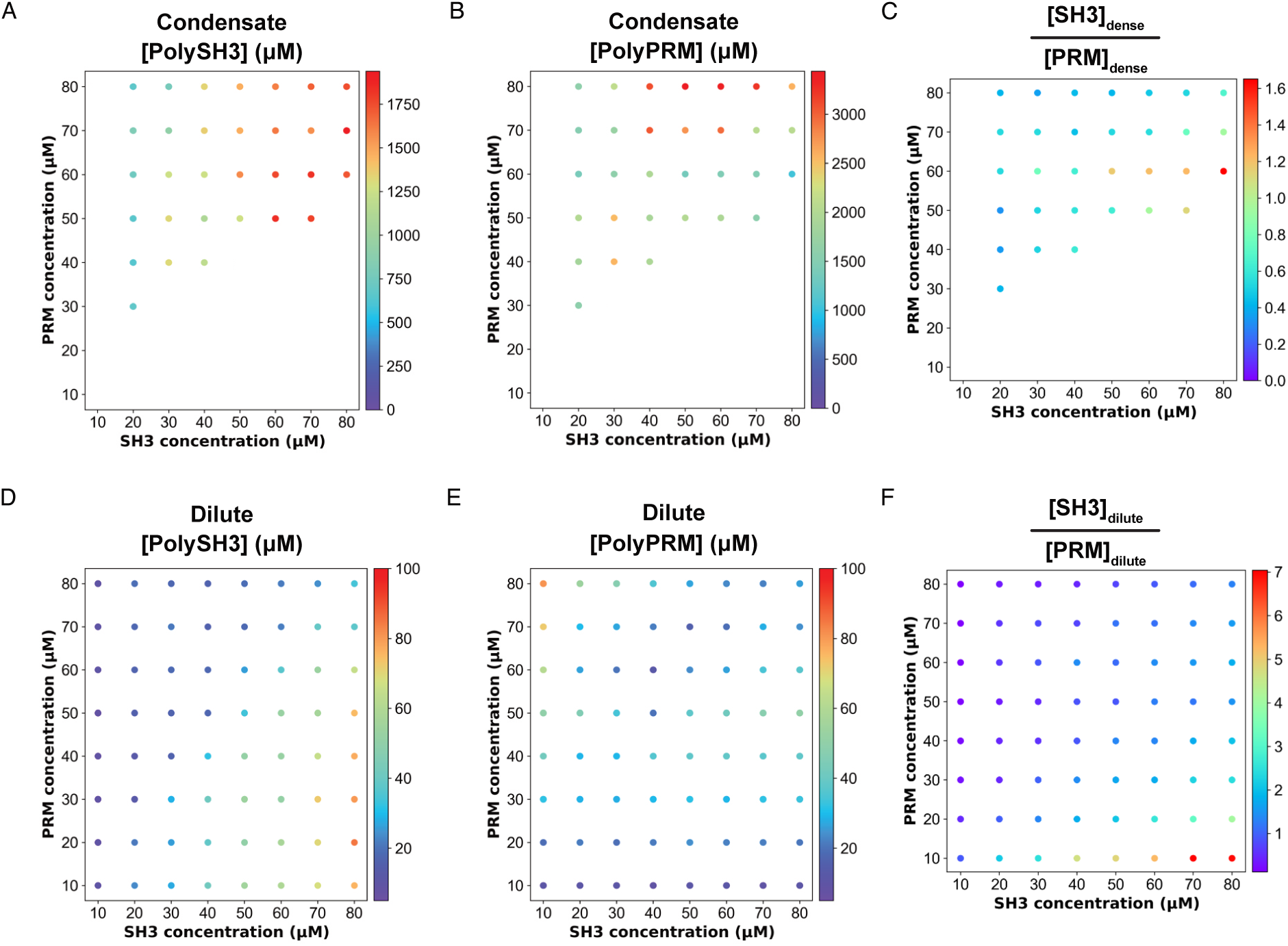
**A)** Experimentally measured dense phase concentrations of polySH3 across the phase diagram. **B)** Experimentally measured dense phase concentrations of polyPRM across the phase diagram. **C)** The ratio of polySH3: polyPRM in the dense phase across the experimental phase diagram. **D)** Experimentally measured dilute phase concentrations of polySH3 across the phase diagram. **E)** Experimentally measured dilute phase concentrations of polyPRM across the phase diagram. **F)** The ratio of polySH3: polyPRM in the dilute phase across the experimental phase diagram.

**Figure S4.**
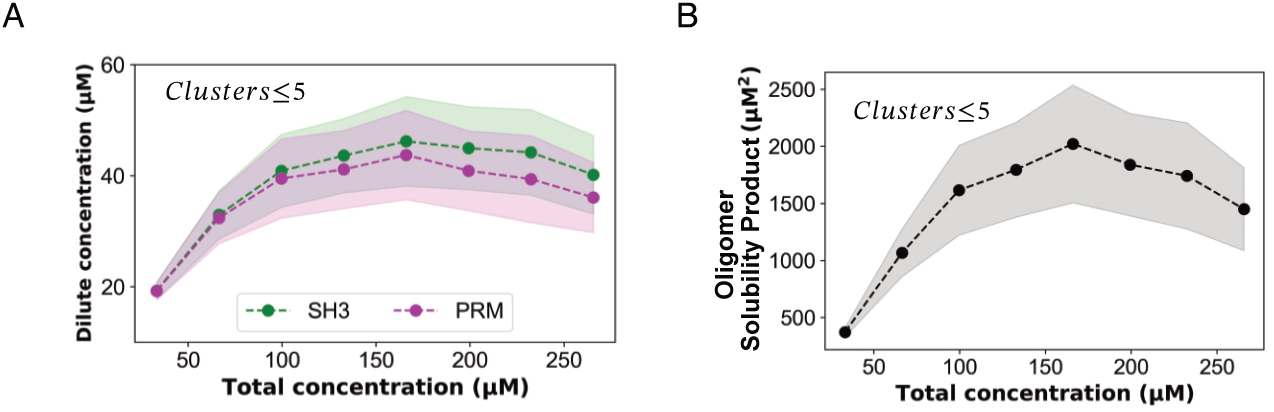
**A)** Simulated concentrations of polySH3 (green) and polyPRM (magenta) along the concentration diagonal when clusters containing five or fewer molecules are quantified. Simulations are performed with heterotypic polySH3 - polyPRM and homotypic polyPRM – polyPRM interactions. **B)** Simulated SP_oligomer_ from simulations performed with heterotypic polySH3 - polyPRM and homotypic polyPRM – polyPRM interactions.

